# pH Effect on the Dynamics of SARS-CoV-2 Main Protease (M^*pro*^)

**DOI:** 10.1101/2020.11.30.404384

**Authors:** Shilpa Sharma, Shashank Deep

## Abstract

The SARS-CoV-2 main protease (M^*pro*^) is a crucial enzyme responsible for the maturation of novel coronavirus, thus it serves as an excellent target for drug discovery. SARS-CoV-2 is found to have similarity with SARS-CoV, which showed conformational changes upon varying pH. There is no study till date on how pH change affect the conformtional flexibilty of SARS-CoV-2 M^*pro*^, therefore, we attempt to find the effect of pH variation through constant pH molecular dynamics simulation studies. Protein is found to be most stable at neutral pH and as pH turns basic protein structure becomes most destabilized. Acidic pH also tends to change the structural properties of M^*pro*^. Our study provides evidence that the flexibility of M^*pro*^ is pH dependent like SARS-CoV M^*pro*^.

## 1 Introduction

COVID-19 is a severe respiratory illness caused due to novel coronavirus, SARS-CoV-2 (severe acute respiratory syndrome coronavirus 2).^1^ Coronaviruses belong to Nidovirales family, these are enveloped, positive-sense RNA viruses, having very long genome around 30000 bases.^2^ 10 kilobases are used for structure and accessory functions and 20 kilobases encode for non-structural proteins.^2^ The replicase gene encodes for the two overlapping polyproteins pp1a and pp1ab, which are critical for replication and transcription. These polyproteins are processed extensively by virus proteinases and the functional proteins are released for carrying out the virus replication.^3^ M^*pro*^ also referred as 3CI^*pro*^, is a 33 kDa cysteine protease responsible for cleaving and processing of viral polyproteins. The structure of SARS-CoV-2 M^*pro*^ free enzyme is recently solved by Zhou et al^4^ at pH 7. M^*pro*^ is a homodimeric protein, shown in Fig. 1. Structure of each monomer is comprised of three domains. Residues 10-99, 100-182 and 198-303 are involved in the formation of domains 1,2 and 3, respectively. Dimerization of M^*pro*^ is regulated by domain 3. Catalytic dyad of the enzyme is formed by residues His 41 and Cys145. Dimerization is necessary for the catalytic activity of M^*pro*^.^5,6^ Due to its role in viral transcription and replication, M^*pro*^ serves as an attractive target for drug designing. Many studies have been carried out to identify the potential inhibitors of M^*pro*^.^7,8^

**Figure 1:**
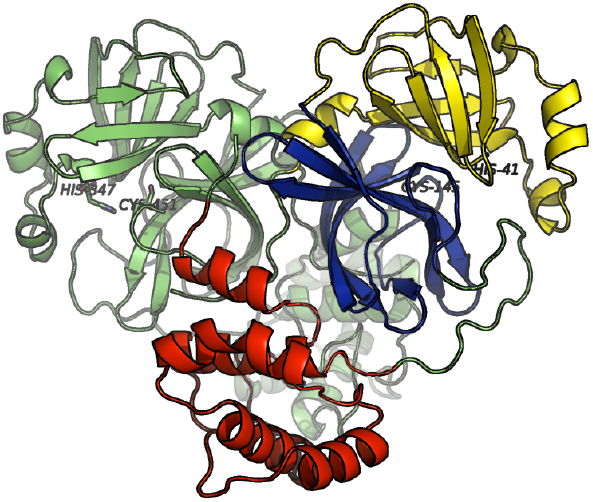
Structure of main protease (M^*pro*^), three domains of monomeric unit are highlighted as yellow, blue and red, respectively.

Function of a protein is closely related to its structure. As structure perturbs from native conformation, its activity changes and under drastic conditions protein may become functionally inactive. Various extraneous conditions like heat, concentration of like and unlike molecules around and pH of surrounding etc affect the protein structure, therefore, their effects are important to be examined on the protein of interest. Proteins are composed of various amino acids having acidic, basic, aliphatic, polar and non-polar side chains. As pH of the surroundings is changed, the protonation states of these side chains also modify which in turn alters several interactions and give protein a distinct structure. Effect of pH was seen on the structure of SARS-CoV M^*pro*^ by Jinzhi et al.,^9^ their study showed that the conformational flexibility of M^*pro*^ is pH dependent. Henedersen et al.^10^ utilized constant pH MD simulations to assess the proton coupled dynamics of various PLpros including SARS-CoV-2. In our knowledge, there is no pH dependant study of SARS-CoV-2 M^*pro*^ structure has been conducted, therefore, in our study, we have seen the effect of pH alteration, ranging from acidic to basic pH, on the structure of SARS-CoV-2 main protease (M^*pro*^) through constant pH molecular dynamics simulations (CpHMD).

## 2 Simulation Details

All atom constant pH molecular dynamics (CpHMD) simulations^11,12^ were performed using AMBER18^13^ and amberff10^14^ force field was used for deriving the bonded and nonbonded parameters. Initial structure of protein was obtained from RCSB (PDB: 6y2e^4^) and protein was solvated using TIP3P^15^ water molecules inside trucated octahedron box of 18 nm. Charges on the protein were neutralized by adding counter ions. Energy minimization was carried out using 1000 steps of steepest descent followed by 4000 steps of conjugate gradient method, keeping backbone atoms restrained. Temperature of the system was slowly increased from 10 K to 300 K over 400 ps under NVT ensemble, followed by equilibration at constant pressure for 4 ns. Berendsen barostat^16^ was used for maintaining constant pressure condition. The study utilized periodic boundary conditions and PME summation^17^ for electrostatic calculations.The shake methodology^18^ was applied to restrict covalently bonded hydrogen atoms. Time step of 2 femtosecond with 8 A^0^ cut off for non-bonded interactions were applied. Convergence of energy, temperature and density were monitored. Parallel to the molecular dynamics simulation, the constant pH method applied in pmemd through Monte Carlo sampling of the Boltzmann distribution of individual protonation states. Distribution of protonation states is influenced by solvent pH, which is set as an external parameter. A total of five CpHMD simulations were run at pH 4, 5, 6, 7 and 8, by varying the solvent pH. All the acidic aminoacids (ASP and GLU) are assumed to be deprotonated and basic aminoacids (ARG, LYS and CYS) to be protonated in this pH range. Protonation states of catalytic HIS 41 and CYS 145 were unchanged, all other twelve histidines present in the protein were set as titrable residues and their protonation states are subjected to change by changing the partial charges on the atoms of titrable residues. Production run was carried out for 100 ns, during production after every 100 steps protonation state change of every titrable residue was attempted. GB salt concentration was set to 0.1 M.

Structural changes were seen through root mean square deviation (RMSD) of backbone atoms, which was calculated with respect to the first frame structure. Residue wise fluctuations were seen through root mean square fluctuation (RMSF) which is time average RMSD. Surface area of protein was calculated using LCPO algorithm by Weiser et al.^19^ For calculating the native contacts, 7 A^0^ cut off was set, active contacts were found to be present if the distance between two C-alpha atoms were within 7 A^0^ at any given time frame. Clustering analysis was done using average linkage method, by keeping epsilon equal to 1. The most populated cluster was selected for comparing substrate binding pockets at different pH. Distance between two residues was calculated using distance tool of cpptraj.^20^

## 3 Results and Discussion

### 3.1 Root Mean Square Deviation

The change in protonation states bring about local fluctuations in protein which then contribute to overall dynamical changes in protein, therefore to see these changes, root mean square deviation (RMSD) was monitored throghout the simulation, as shown in Fig. 2(a). Initially, RMSD values increased slowly at all the pH values but a sudden increase in RMSD was observed at pH 8 around 50 ns (1 ns=500 frames), which kept on increasing till 80 ns and became constant after that. For a better understanding, probability distribution curve of each RMSD value was constructed, (Fig. 2(b)), two peaks were observed around 1.8 A^0^, 2.3 A^0^ at pH 4 and around 2.4 A^0^, 2.9 A^0^ at pH 8. For pH 5, 6 and 7 RMSD values having maximum prob-ablities were observed at 2.2 A^0^, 1.9 A^0^ and 2 A^0^, respectively. From RMSD analysis, it can be seen that at pH 4 and pH 8 protein sampled large number of conformations and fluctuations were maximum at pH 8, minimum at pH 7.

**Figure 2:**
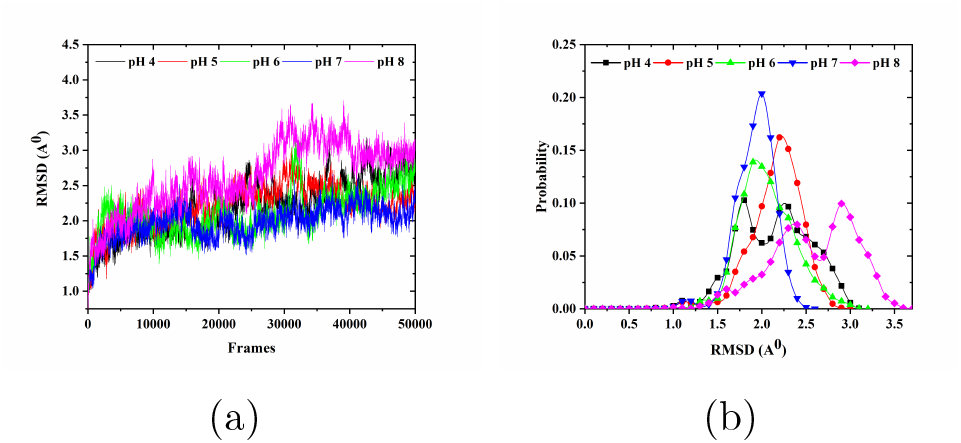
(a) Backbone root mean square deviation plot and (b) probability distribution of RMSD values at various pH.

### 3.2 Root Mean Square Fluctuation

Contribution of each residue towards the total structural deviation was seen through root mean square fluctuation (RMSF) analysis, Fig. 3(a)–3(b). At pH 4, larger fluctuations were seen for residues 117-123, 141-143, 196-199, 209-216, 226-263, 280-288 and 293-301 in chain A, comparatively smaller fluctuations were seen in chain B. At pH 5, residues 50-52, 72, 96-97, 184-187 and 299 of chain A and residues 1, 45-51, 140, 155, 167-170, 189-196, 216-218, 270-274, 304-306 of chain B showed more changes. At pH 6, residue 1-3, 59-65, 109-112, 128-129, 132, 138-139, 154-156, 195-203, 213-215, 218, 220-249, 257-258, 262-263, 266, 269, 273, 290-301 of chain A and residues 1-3, 58-65, 118-120, 168-169, 244-258 and 305-306 of chain B were shown to be more fluctuating. Fluctuations were found be minimum in both the chains at pH 7 as compared to that of at other pH values. At pH 8, chain A showed relatively lesser fluctuations as compared to chain B. In chain A, residues 1-2, 52-53, 138, 183-185, 192, 194, 274 and 19-27, 39-90, 116-122, 137-146, 167-172, 176-180, 183, 191-192, 226-229, 231-233 and 275-276 residues in chain B were largely destabilized. At pH 4, mainly domain 3 of chain A was destabilized with highest RMSF values. As pH increases from 4 to 5, fluctuations were also observed in domains 1 and 3. At pH 6, destabilization in all three domains became dominant and increased even more at pH 8. Fluctuations in domain 1 and 2 affects substrate binding to M^*pro*^ and in domain 3 affect dimerization. Above observations suggest that all the important domains of M^*pro*^ namely, 1,2 and 3 gets destabilized as pH becomes acidic or basic.

**Figure 3:**
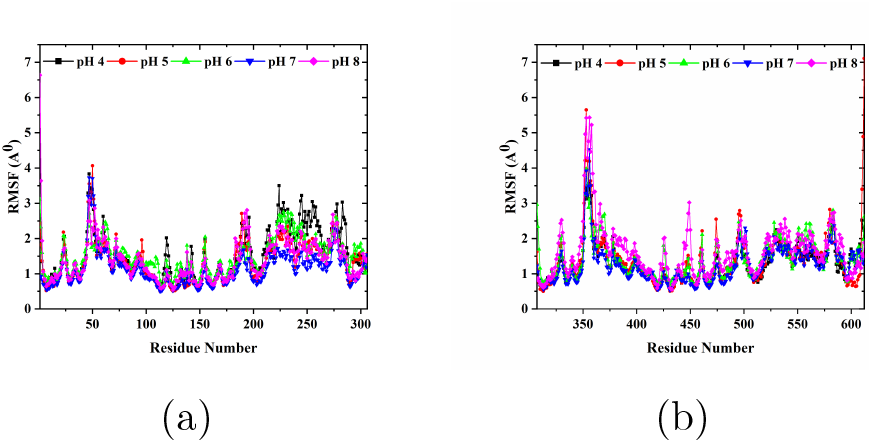
(a) Backbone root mean square fluctuation plot for chain A and (b) for chain B at various pH.

### 3.3 Solvent Accessible Surface Area and Native Contacts

Solvent accessible surface area (SASA) and number of native contacts can be used to monitor the proper folding of protein, therefore, these were calculated at all the pH (Fig. 4). At pH 4, 5 and 7, only a small change in SASA was observed with time, most probable values of SASA were equal to 267.37 nm^2^, 269.31 nm^2^ and 261.98^2^, respectively. At pH 6, SASA started increasing around 20 ns and kept on increasing till 100 ns, with two peaks at 264.27 and 275 nm^2^ in probability plot. At pH 8, value of SASA first decreased and then started increasing again after 20 ns, a wide probability distribution of SASA was obtained with 270.78 nm^2^ having maximum probability.

**Figure 4:**
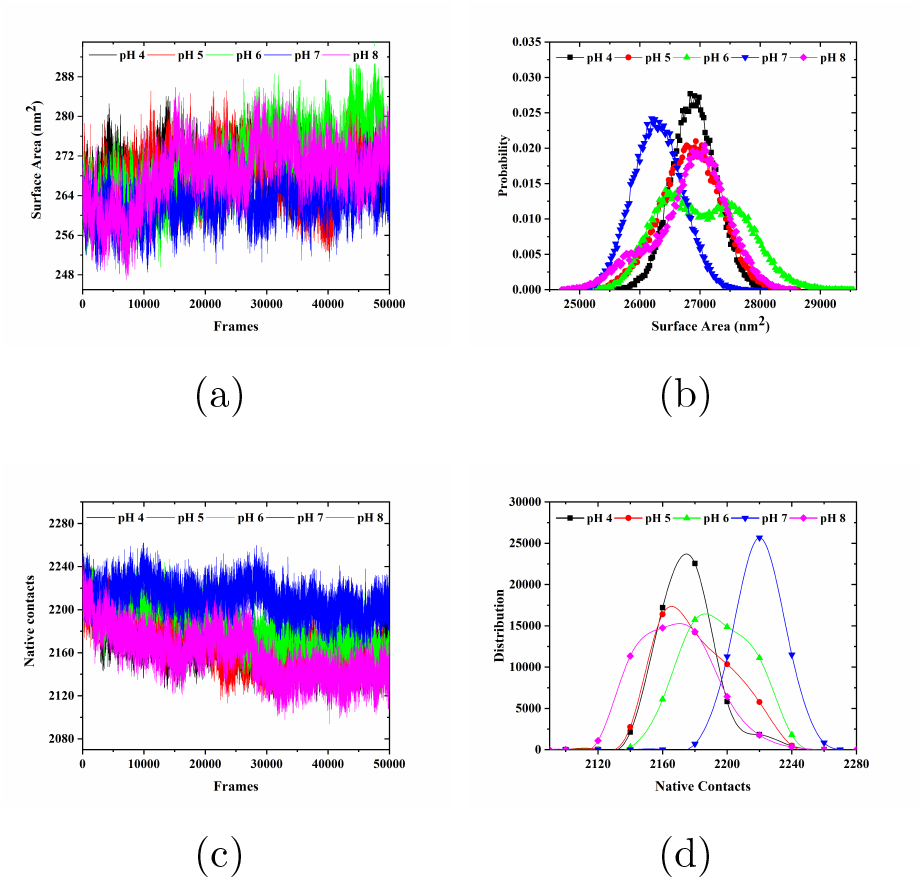
Plots for (a) Accessible surface area (ASA) and (b) probability distribution of ASA values, (c) native contacts in protein, (d) probability distribution of native contacts' values at different pH.

In the beginning, native contacts were around 2240 at all pH, after few nanoseconds native contacts started decreasing and at pH 8 maximum decrease in contacts was observed, Fig. 4(c). At pH 7, only slight changes in native contacts were observed. From probability distribution curve, native contacts were found to be highest, 2220, at pH 7. At pH 4, 2170 native contacts were present. At pH 5, 6 and 8, a broad distribution of native contacts was seen, with increase in contacts from pH 5 to 6 and least number of native contacts were there at pH 8, Fig. 4(d). Surface area was found to be minimum and native contacts were found to be maximum at pH7, implying that the structure of M^*pro*^ is most stable at neutral pH.

### 3.4 Distance Analysis

Dimerization of main protease is essential for its activity (because the N-finger of each of the two protomers interacts with Glu166 of the other protomer and thereby helps shape the S1 pocket of substrate binding site), its monomeric state is not active. Substrate binding site of enzyme is shaped through the interaction of N-finger of one monomer and Glu166 of another monomer, therefore, distance between residues Ser1 and Glu166 was monitored to ensure intact dimer formation throughout the simulation at all pH values, Fig. 5(a). At pH 7, distance between Ser1 and Glu166 was remained constant at all time frames. At pH 4, 5 and 6, this distance increased with time and value fluctuated between 8 A^0^ and 12 A^0^. At pH 8, distance started increasing after 20 ns and a sudden increase was observed around 45 ns followed by a constant fluctuation around 16 A^0^ at all frames. Similarly, in probability histogram multiple peaks are observed at pH 4, 5, 6 and 8, however, a single peak was observed at pH 7, Fig. 5(b), which suggests that dimerization process is weak at acidic and basic pH.

**Figure 5:**
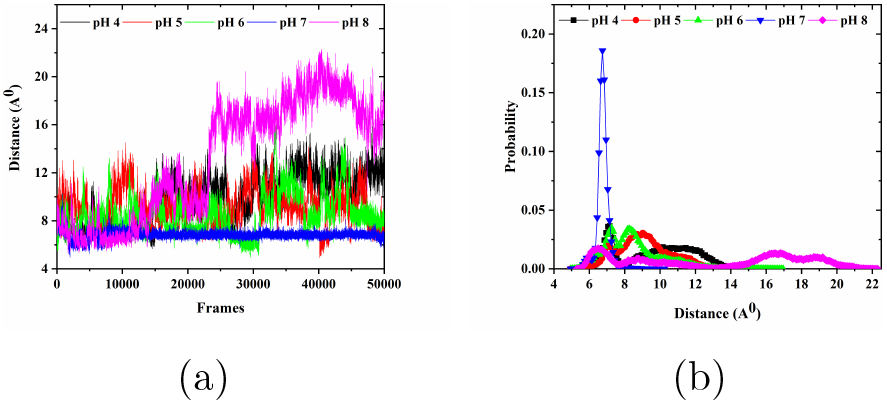
Plots for (a) distance between N-finger of chain A and Glu166 of chain B, (b) probability distribution of distance at every pH value.

### 3.5 Clustering Analysis and Orientation of Substrate Binding Pocket

From clustering analysis, representative structures of the most populated clusters were obtained at all pH values. Proper orientation of active site residues is necessary for protein to carry out its function. The active sites, of the obtained structures of M^*pro*^ from clustering, were compared, Fig. 6(a)–6(e). His 41 and Cys 145 forms the catalytic dyad of protein, their proper orientation was observed only at pH 7, Fig. 6(d). At all the pH other than pH 7, N^ϵ2^ of His 41 was not found to be orienting towards S^γ^ of Cys 145, as shown in Fig. 6(a), 6(b), 6(c) and 6(e). Thus, the orientation of catalytic dyad of M^*pro*^ S1 binding pocket also gets affected by pH.

**Figure 6:**
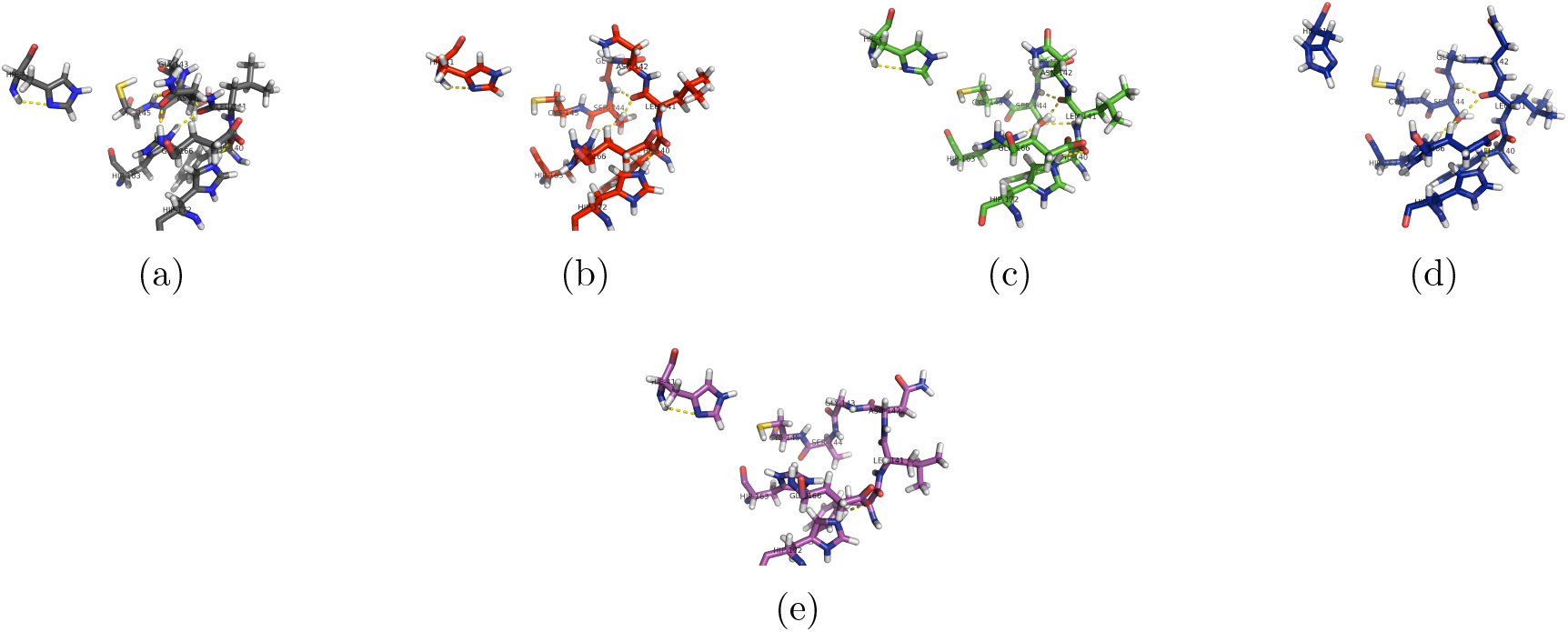
S1 binding pocket of M^*pro*^ at (a) pH4, (b) pH5, (c) pH6, (d) pH7 and (e) pH 8.

## 4 Conclusion

Our study has demonstrated that structure of SARS-CoV-2 main protease (M^*pro*^) shows pH dependence. At basic pH (pH 8), the structure of M^*pro*^ was found to be least stable and as pH becomes more acidic, structure of M^*pro*^ gets destabilized.The structure of M^*pro*^ showed maximum stability at neutral pH. At all the pH other than neutral pH, dimerization and S1 binding pocket were found to be affected. Our study throws light on how an environmental factor like pH can affect the integrity of active site of M^*pro*^ and encourage researchers to explore the small molecules that can modulate the intracellular pH to prevent the replication by lowering the activity of M^*pro*^.

## 5 Acknowledgment

SS thanks UGC India for fellowship. Authors thank the IIT Delhi HPC facility for computational resources. Authors also thank Scf-Bio for providing the access of AMBER18.

## References

(1) Subbarao, K.; Mahanty, S. Respiratory Virus Infections: Understanding COVID-19. Immunity 2020, 52, 905–909.

(2) Fehr, A. R.; Perlman, S. Coronaviruses: an overview of their replication and pathogenesis. Methods in molecular biology (Clifton, N.J.) 2015, 1282, 1–23.

(3) Ulferts, R.; Imbert, I.; Canard, B.; Ziebuhr, J. Expression and Functions of SARS Coronavirus Replicative Proteins. Molecular Biology of the SARS-Coronavirus 2009, 75–98.

(4) Zhang, L.; Lin, D.; Sun, X.; Curth, U.; Drosten, C.; Sauerhering, L.; Becker, S.; Rox, K.; Hilgenfeld, R. Crystal structure of SARS-CoV-2 main protease provides a basis for design of improved-ketoamide inhibitors. Science 2020, 368, 409–412.

(5) Anand, K.; Ziebuhr, J.; Wadhwani, P.; Mesters, J. R.; Hilgenfeld, R. Coronavirus Main Proteinase (3CLpro) Structure: Basis for Design of Anti-SARS Drugs. Science 2003, 300, 1763–1767.

(6) Jin, X., Zhenmingnad Du; Xu, Y.; Deng, Y.; Liu, M.; Zhao, Y.; Zhang, B.; Li, X.; Zhang, L.; Peng, C.; Duan, Y. et al. Structure of Mpro from SARS-CoV-2 and discovery of its inhibitors. Nature 2020, 582, 289–293.

(7) Sharma, S.; Deep, S. In-silico drug repurposing for targeting SARS-CoV-2 main protease (Mpro). Journal of Biomolecular Structure and Dynamics 2020, 0, 1–8, PMID: 33179568.

(8) Maffucci, I.; Contini, A. In Silico Drug Repurposing for SARS-CoV-2 Main Proteinase and Spike Proteins. Journal of Proteome Research 2020, 19, 4637–4648, PMID: 32893632.

(9) Tan, J.; Verschueren, K. H. G.; Anand, K.; Shen, J.; Yang, M.; Xu, Y.; Rao, Z.; Bigalke, J.; Heisen, B.; Mesters, J. R. et al. pH-dependent conformational flexibility of the SARS-CoV main proteinase (M(pro)) dimer: molecular dynamics simulations and multiple X-ray structure analyses. J Mol Biol. 2005, 354, 25–40.

(10) Henderson, J. A.; Verma, N.; Harris, R. C.; Liu, R.; Shen, J. Assessment of proton-coupled conformational dynamics of SARS and MERS coronavirus papainlike proteases: Implication for designing broad-spectrum antiviral inhibitors. The Journal of Chemical Physics 2020, 153, 115101.

(11) Swails, J. M.; York, D. M.; Roitberg, A. E. Constant pH Replica Exchange Molecular Dynamics in Explicit Solvent Using Discrete Protonation States: Implementation, Testing, and Validation. Journal of Chemical Theory and Computation 2014, 10, 1341–1352, PMID: 24803862.

(12) Mongan, J.; Case, D. A.; McCammon, J. A. Constant pH molecular dynamics in generalized Born implicit solvent. Journal of Computational Chemistry 2004, 25, 2038–2048.

(13) Case, D. A.; Cheatham III, T. E.; Darden, T.; Gohlke, H.; Luo, R.; Merz Jr., K. M.; Onufriev, A.; Simmerling, C.; Wang, B.; Woods, R. J. The Amber biomolecular simulation programs. Journal of Computational Chemistry 2005, 26, 1668–1688.

(14) Wang, J.; Cieplak, P.; Kollman, P. A. How well does a restrained electrostatic potential (RESP) model perform in calculating conformational energies of organic and biological molecules? Journal of Computational Chemistry 2000, 21, 1049–1074.

(15) Jorgensen, W. L.; Chandrasekhar, J.; Madura, J. D.; Impey, R. W.; Klein, M. L. Comparison of simple potential functions for simulating liquid water. The Journal of Chemical Physics 1983, 79, 926–935.

(16) Berendsen, H. J. C.; Postma, J. P. M.; van Gunsteren, W. F.; DiNola, A.; Haak, J. R. Molecular dynamics with coupling to an external bath. The Journal of Chemical Physics 1984, 81, 3684–3690.

(17) Petersen, H. G. Accuracy and efficiency of the particle mesh Ewald method. The Journal of Chemical Physics 1995, 103, 3668–3679.

(18) Krutler, V.; van Gunsteren, W. F.; Hnenberger, P. H. A fast SHAKE algorithm to solve distance constraint equations for small molecules in molecular dynamics simulations. Journal of Computational Chemistry 2001, 22, 501–508.

(19) Weiser, J.; Shenkin, P. S.; Still, W. C. Approximate atomic surfaces from linear combinations of pairwise overlaps (LCPO). Journal of Computational Chemistry 1999, 20, 217–230.

(20) Roe, D. R.; Cheatham, T. E. PTRAJ and CPPTRAJ: Software for Processing and Analysis of Molecular Dynamics Trajectory Data. Journal of Chemical Theory and Computation 2013, 9, 3084–3095, PMID: 26583988.

